# Spatial structure, chemotaxis and quorum sensing shape biomass accumulation in complex systems

**DOI:** 10.1101/2023.03.28.534616

**Authors:** David Scheidweiler, Ankur Deep Bordoloi, Vladimir Sentchilo, Monica Bollani, Philipp Engel, Pietro de Anna

## Abstract

Biological tissues, sediments, or engineered systems are spatially structured media with a tortuous and porous structure that host the flow of fluids. Such complex environments can influence the spatial and temporal colonization patterns of bacteria by controlling the transport of individual bacterial cells, the availability of resources, and the distribution of chemical signals for communication. Yet, due to the multi-scale structure of these complex systems, it is hard to assess how different biotic and abiotic properties work together to control the accumulation of bacterial biomass. Here, we explore how flow mediated interactions allow the gut commensal Escherichia coli to colonize a porous structure that is composed of heterogenous dead-end pores (DEPs) and connecting percolating channels, i.e. transmitting pores (TPs), mimicking the structured surface of mammalian guts. We find that in presence of flow, gradients of the quorum sensing (QS) signaling molecule autoinducer-2 (AI-2) promote E. coli chemotactic accumulation in the DEPs. In this crowded environment, the combination of growth and cell-to-cell collision favors the development of suspended bacterial aggregates. This results in hot-spots of resource consumption, which, upon resource limitation, triggers the mechanical evasion of biomass from glucose and oxygen depleted DEPs. Our findings demonstrate that microscale medium structure and complex flow coupled with bacterial quorum sensing and chemotaxis control the heterogenous accumulation of bacterial biomass in a spatially structured environment, such as villi and crypts in the gut or in tortuous pores within soil and filters.

## Introduction

Bacteria typically inhabit spatially complex environments, ranging from biological tissues such as the gut, to soil and sediments, where resources, chemical stimuli, and physical constraints are patchy and transient [1–3]. In these environments, fluids are forced to flow within pore spaces confined between irregular surfaces. Several features characterize such confined systems, including their material composition [4,5], pore-size, pore-morphology, and permeability to fluids [6]. Spatial variability (referred to here as, heterogeneity) in these properties appears in most realistic scenarios. The gut environment, for instance, is known for its complex spatial heterogeneity, ranging across multiple scales - from a few microns (e.g. villi and crypts) to several millimeters (e.g. circular folds) and meters (small to large intestine), leading to complex transport of fluid, chemical gradients, and spatial organization of microbial communities [2]. Yet, how the heterogeneity of such environments determines bacterial colonization, by controlling the transport of individual bacteria [7,8], cell-to-cell interactions [9], and growth [10], remains poorly understood.

Recent research suggests that the spatial arrangement of bacteria in the gut is a critical factor influencing their function and interaction with the host [2,11,12]. Recent studies have demonstrated that the gut microbiota exhibits intricate spatial patterns, with distinct bacterial species occupying specific niches within the gut lumen and mucosal surface (i.e. villi and crypts) [13,14]. These patterns are influenced by various factors, including flow dynamics, nutrient availability, and ecological interactions among bacterial species [5,15]. However, studying the impact of these factors in vivo comes with obvious constraints. Therefore, researchers adopted alternative strategies such as microfluidics [16,17], organoids [18,19], and in-silico approaches [20,21] to study the impact of flow on gut related microorganisms. The results of these studies indicate that fluid movement in the gut can significantly affect bacterial behavior, including their spatial organization and interactions with the host.

Structural properties of heterogenous environments control the flow of fluids (i.e. advection) [22,23], transport of suspensions (e.g. bacteria) [7], solutes (e.g. nutrients) and the establishment of their gradients [24]. By combining sensory receptors and motility systems, bacteria direct their motion towards or away from gradients in dissolved oxygen, carbon sources or self-secreted metabolites [25–27]. This behavior, known as chemotaxis, entails a population benefit for bacteria at the cost of resources invested in motility [8,28,29]. Earlier research showed that chemotactic motility influences bacterial dispersal in presence of stationary [19–21], and dynamic gradient conditions [10]. However, in both scenarios, the effects of chemotaxis are limited to relatively short periods of time (up to a few tens of minutes). After an initial transient period, molecular diffusion dissipates the chemical gradient, terminating the chemotactic activities even when the chemoattractant is a self-secreted metabolite [30,31]. Therefore, observations on persistent chemotactic gradients, leading to cell accumulations over several hours - a time scale comparable to the one of bacterial colony growth - have never been reported.

Bacteria can sense concentration and gradients of self-secreted metabolites. This sensory ability allows them to detect differences in population density and coordinate gene regulation: a mechanism called quorum sensing (QS) [32]. QS is initiated by the secretion of signaling molecules, known as AutoInducers (AI), which may accumulate in an extracellular environment with increasing cell density. This cell-to-cell “communication” mechanism enables bacteria to modulate collective behaviors, such as secretion of virulence factors, motility, and biofilm formation [33–36]. The latter leads to the transition of bacteria from the planktonic state to multicellular architectures embedded within a matrix of extracellular polymeric substances (EPS) [37,38] that allow them to persist in diverse environments [2,13]. While certain signaling systems are species-specific (known as QS type-1; such as for *Vibrio harveyi*), other signaling systems enable interspecies communication (known as QS type-2) [33]. This is the case of the mammalian gut commensal *Escherichia coli*, whose cells produce, detect and uptake interspecies signaling molecule AutoInducer-2 (AI-2). QS type-2 allows different species to sense their mutual abundance as well as interfere with neighboring cells [33]. Additionally, the regulation of QS type-2 can be affected by different resources availability and cellular metabolism. In presence of glucose, the carbon catabolite repression inhibits the AI-2 uptake of *E. coli*, therefore hindering QS induction [34,39,40].

The physical structure of the surrounding system controls the spatial and temporal availability of resources and AIs by modulating its local transport [9]. On the one hand, flow can disrupt QS by displacing the produced signaling molecules. On the other hand, stagnant regions promote the AIs accumulation, controlling the local QS induction, as shown for of *Staphylococcus aureus* and *Vibrio cholerae* [41]. Moreover, different studies demonstrated that *E. coli*, can sense AI-2 through the Tsr chemoreceptor of L-Serine, and the regulator of the flagellar motor CheY, allowing cells to swim towards gradients of the signaling molecules themselves [30,31]. These findings suggest that the interplay between physical structure, flow, and sensing regulation, might have remarkable implications on the bacterial colonization in complex environments.

In this study, we investigate how the interplay of the spatial structure, fluid-flow, chemotaxis and QS controls the colonization and biomass accumulation within a porous micro-environment. To this end, we use a microfluidic model featuring a structure that comprise Dead-End Pores (DEPs) connected to a network of Transmitting Pores (TPs) [23], typical of spatially complex systems, such as biological tissues or soil. We show that, in such complex structures, QS is used by *E. coli* to turn stagnant DEPs into hot-spots of bacterial colonization. We demonstrate three primary mechanisms that control the overall colonization process. During the early times, accumulation of individual cells within the DEPs is driven by chemotaxis towards self-produced AI-2. Then, cell aggregation in DEPs is further promoted by motility towards gradients of AI-2 released by bacterial clusters and cell-to-cell collision. At late times, resources limitation and accumulation of AI-2 lead to biofilm formation induced by QS, which mechanically force daughter cells out of resource limited zones. These observations are justified by the persistence of self-produced AI-2 gradients for long times, tens of hours, at the DEP-TP interface. Our results indicate that the spatial organization of complex structures plays a pivotal role in local colonization of *E. coli*.

## Results

### A microfluidic set-up to study *E. coli* colonization of a heterogeneous porous system

Within complex physical structures, the hydrodynamic competition between advection dominated pores (controlled by fluid displacement) and stagnant zones (controlled by molecular diffusion) results in heterogeneous transport conditions for solutes and microorganisms. To explore the role of microscale structures on the invasion and colonization of the mammalian gut commensal bacterium *Escherichia coli* MG1655, we design a porous micromodel (Fig. 1a,b) comprising cavity-like irregular structures commonly found in the gut (Fig. 1c) [2,42]. With this micromodel as a template, we build Polydimethylsiloxane (PDMS) based microfluidic devices featuring cavity structures (see Fig.1c). These cavities, referred to here as the Dead-End Pores (DEPs), are connected to a network of percolating channels, called the Transmitting Pores (TPs). We segregate the two pore-features for the entire system (see Methods); a representative section is shown in Fig 1d.

**Figure 1.**
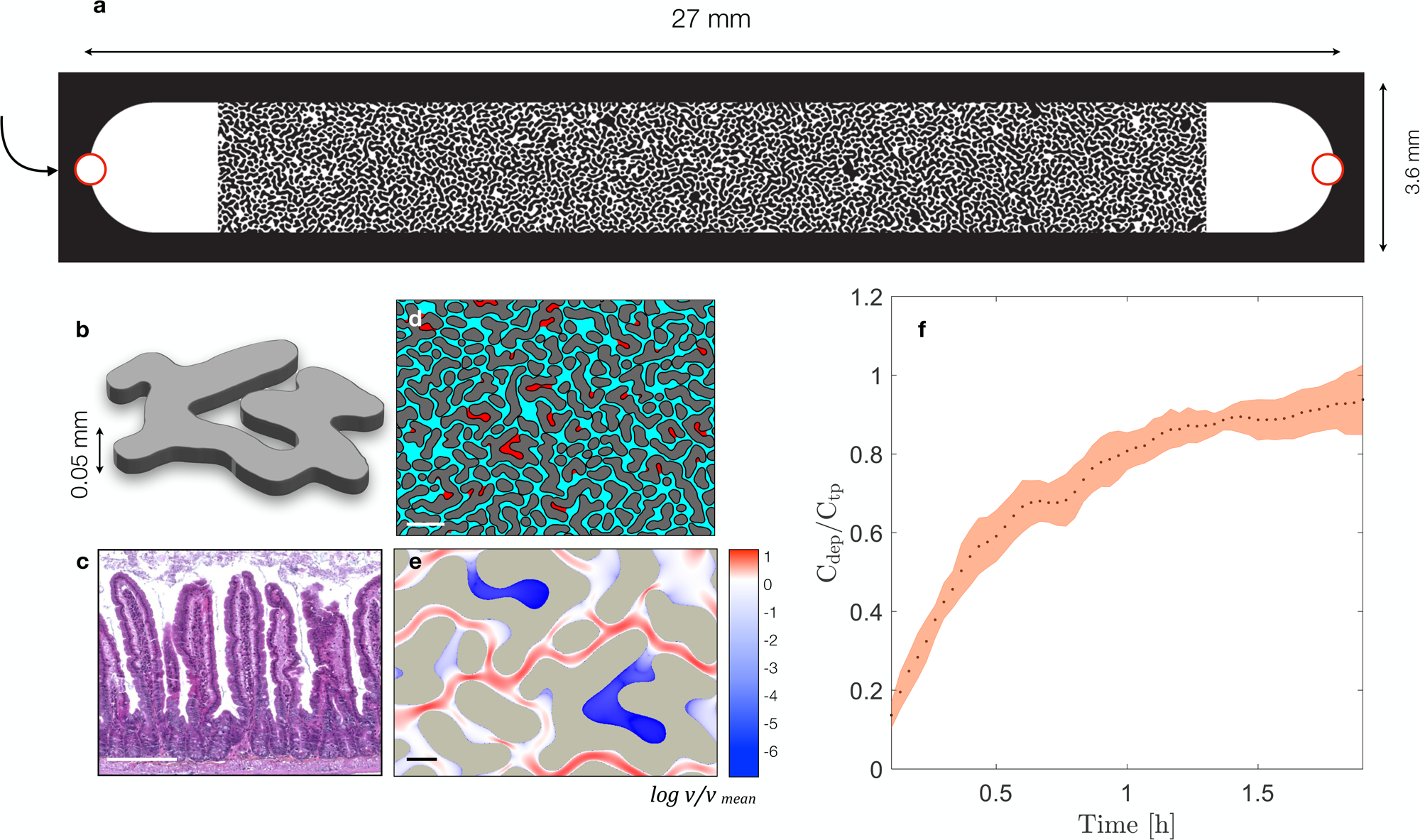
Microfluidic device, geometry, flow and transport characterization. (A) Conceptual diagram of the experimental set-up, (B) highlighting one single impermeable obstacle. (C) Light microscopy image which highlights the complex geometry of mice intestine. Scale bar 200 µm. Courtesy of D. Hardy, Histopathology center, Institut Pasteur. (D) Geometrical discretization into transmitting pores (cyan) and dead-end pores (red). Scale bar 200 µm. (E) Velocity field modulus, normalized by its own averaged value and shown in logarithmic scale, derived from the computed Stokes flow solutions in a subsection of the microfluidic geometry used in the experiments. Scale bar 50 µm. (F) Retention curve representing the ratio between the concentration of bacteria in the dead-end pores (C_dep_) and their concentration in the transmitting pores (C_tp_) at different times.

Prior to our biomass growth measurements, we employ the following set of protocols to homogeneously distribute bacteria across the microfluidic device (see Methods). In these experiments, we use wild-type *E. coli* strain MG1655 (WT) that can produce AI-2, do chemotaxis towards its gradients and do QS. Each experiment is accompanied by an independent experiment wherein the WT strain is replaced by a mutant (*E. coli* Δ*luxS)* lacking the AI-2 synthase *LuxS*. First, we saturate the device with a motility buffer (10 mM potassium phosphate, 0.1 mM EDTA, 10 mM lactate, 1 mM methionine, pH 7.0), followed by a sharp injection (see [23]) of bacterial suspension. At this stage, the bacteria are suspended only within the motility buffer, which allows the injected bacteria to swim (propelled by their flagella), but not to divide and grow. We set the bacteria injection flow rate at Q = 0.1 µL/min, such that the average, Darcy, velocity in the porous medium is comparable to the measured average swimming speed of this *E. coli* strain (Supplementary Fig. S1), and to the mean fluid velocity in human gut (20 µm/sec) [16]. We compute the local fluid velocity within the porous medium by numerically solving the two-dimensional steady state incompressible Stokes flow equations in a subsection of the microfluidics geometry (see Methods). The local fluid velocity spans over several order of magnitude across TPs and DEPs: a portion of the computed velocity field modulus is shown with a logarithmic scale in Fig. 1e. We measure the bacterial swimming speed via a separate experiment under no-flow condition and find it to be identical for the two *E. coli* strains (WT and Δ*luxS* mutant, see Supplementary Fig. S1). With these experimental conditions, it takes approximately 2 hours (corresponding to the injection of 5 times the microfluidics volume) for suspended *E. coli* cells to invade and homogeneously fill the entire pore space. The macroscopic retention curve in Fig. 1f shows that the ratio of cell concentration, averaged over three replicas, within the two pore features DEP and TP (C_dep_/C_tp_) approaches unity at approximately 5 pore volumes (2 hours). The shaded area represents the standard deviation among replicas. Once the medium is homogeneously filled with suspended cells, we switch to the continuous injection of a sterile nutrient solution (see Methods).

### *E. coli* biomass distribution in complex structures

To study the role of transport on chemotaxis and QS through such complex landscape, we examine the overall spatial biomass accumulation of the WT *E. coli* strain MG1655 (WT), contrasted with the mutant experiments (Δ*luxS)*. In each experiment, replicated 3 times, once the bacterial cells homogeneously fill all the pore space, we switch to the continuous injection of a sterile M9 minimal medium solution for 50 hours. We supplement this solution with glucose as the sole carbon source at a concentration comparable with that in mammalian guts (here we used 5 mM) [43]. Through the entire experimental time, as shown in Fig. 2a, the WT strain exhibits a systematically higher accumulation of biomass within the DEPs compared to the TPs. Strikingly, despite DEPs represent only 8% of the total volume, WT cells accumulated five times more within DEPs than in TPs. Overall, as shown in Fig. 2b-c, DEPs contribute to more than 35 ± 1.6% of the total biomass in the system (as quantified with image processing, see Methods). In contrast, while the total biomass of the WT and Δ*luxS* in the system are approximately the same, the biomass of Δ*luxS* accumulated in the DEPs accounts for only 8.1 ± 0.6 % (i.e. mirroring the fraction of the porous medium occupied by the DEPs). As quantitatively captured in Fig. 2b-c, such enhanced cell accumulation clearly demonstrates that the DEPs act as “hot-spots” for microbial accumulation and metabolic activity for WT but not for the Δ*luxS*.

**Figure 2.**
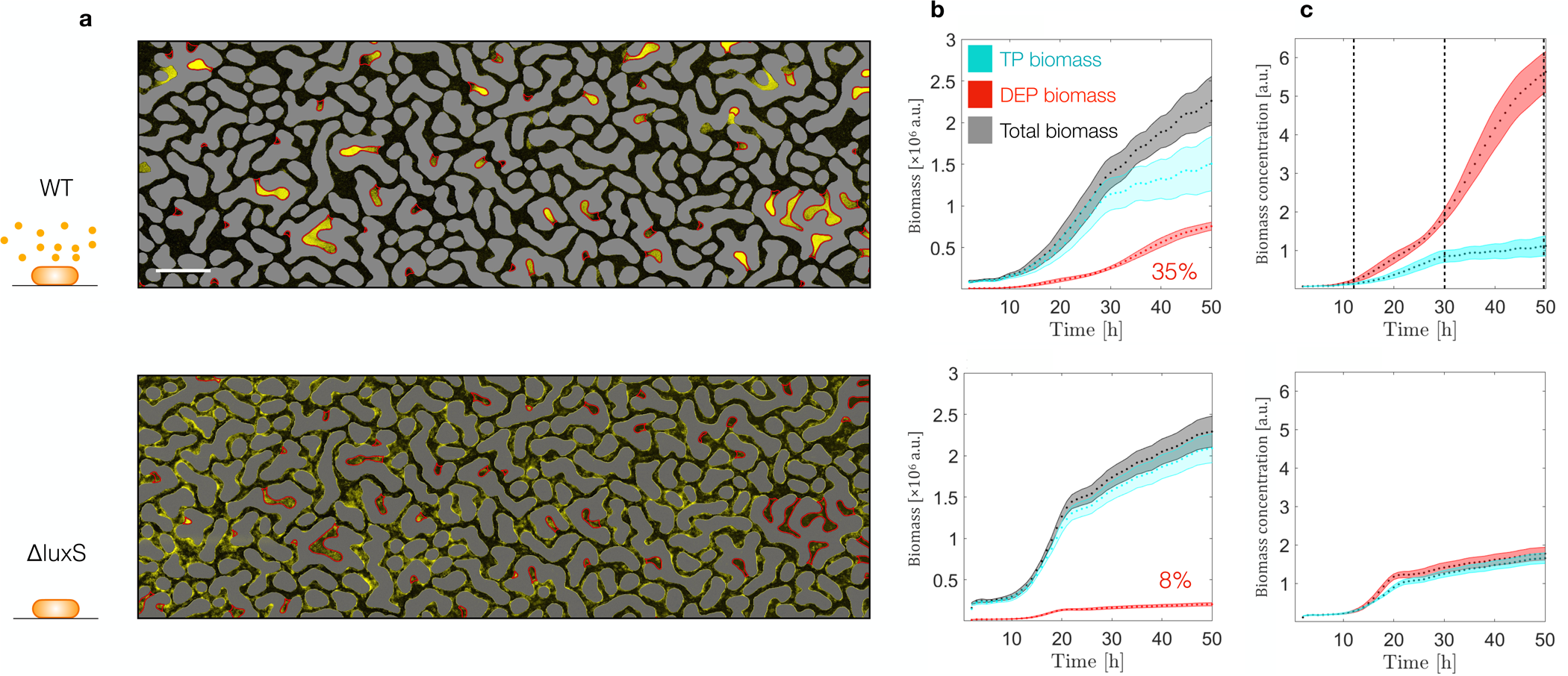
Macroscopic biomass accumulation, for *E. coli* WT (top panels), and *E. coli ΔluxS* (bottom panels). (A) Biomass accumulation through the microfluidic geometry at the end of the experiment (50 hours). Scale bar 200 µm. (B) Dynamics of biomass accumulation in the TP (cyan), in the DEP (red), and in the overall geometry (gray). (C) Dynamics of the biomass concentration in the TP (cyan), in the DEP (red).

Certainly, cells are expected to detach from the TPs due to fluid shear, the average shear rate is *u_m_/ h* = 0.4 *s*^+1^ (*h* begin the chip thickness), where mean fluid velocity is *u_m_*∼20μm/*s*. Both WT and *ΔluxS* strains show similar ability to persist advection-induced shear resulting in comparable bacteria concentrations in the TPs (Fig. 2b and c). Moreover, control experiments injecting higher glucose concentration (50 mM; SI Fig. 2) show that the biomass of WT still accumulates in the DEP, but it reached a higher carrying capacity only TPs, and not in DEPs. These observations suggest that mechanisms other than flow-induced shear promote the higher accumulation of WT biomass in the DEP. We argue that the difference in growth between WT and *ΔluxS* is due to multiple biological activities related to chemotaxis and QS that only the former is capable of. We identify three distinct phases (see dashed lines Fig. 2c) during the bacterial accumulation, characterized by the differences in spatial distributions of biomass: (i) its accumulation in DEPs, (ii) differentiation into aggregates, (iii) biofilm formation. We discuss these phases in further detail to elucidate the underlying biotic and abiotic factors at play.

### Early times - Chemotaxis towards AI-2 promotes biomass accumulation in DEPs

During the first 12h of nutrient injection, we observe that WT cells preferentially accumulate in DEPs (Fig. 3a), resulting in more than 2.5-fold increase in the concentration of suspended cells (Fig 3b). In contrast, *ΔluxS* cells disperse homogeneously across both TPs and DEPs. This discrepancy is not the result of a different motility behavior, as WT and *ΔluxS* show the same velocity distribution (Supplementary Fig S1). Instead, we hypothesize that the higher concentration of WT cells in DEPs is controlled by chemotaxis towards the self-secreted AI-2, which accumulates within the stagnant DEPs, while being advected along TPs. It is important to note that in presence of glucose, WT cells keep producing AI-2 while catabolite repression inhibits AI-2 uptake from the extracellular environment [34,39,40]. Therefore, we expect cells to continuously produce AI-2 until glucose is depleted from the pore space. While advection removes the AI-2 molecules from the TPs, they accumulate in the DEPs leading to AI-2 gradients at the entrance of each DEP that triggers chemotactic migration of WT towards the DEPs.

**Figure 3.**
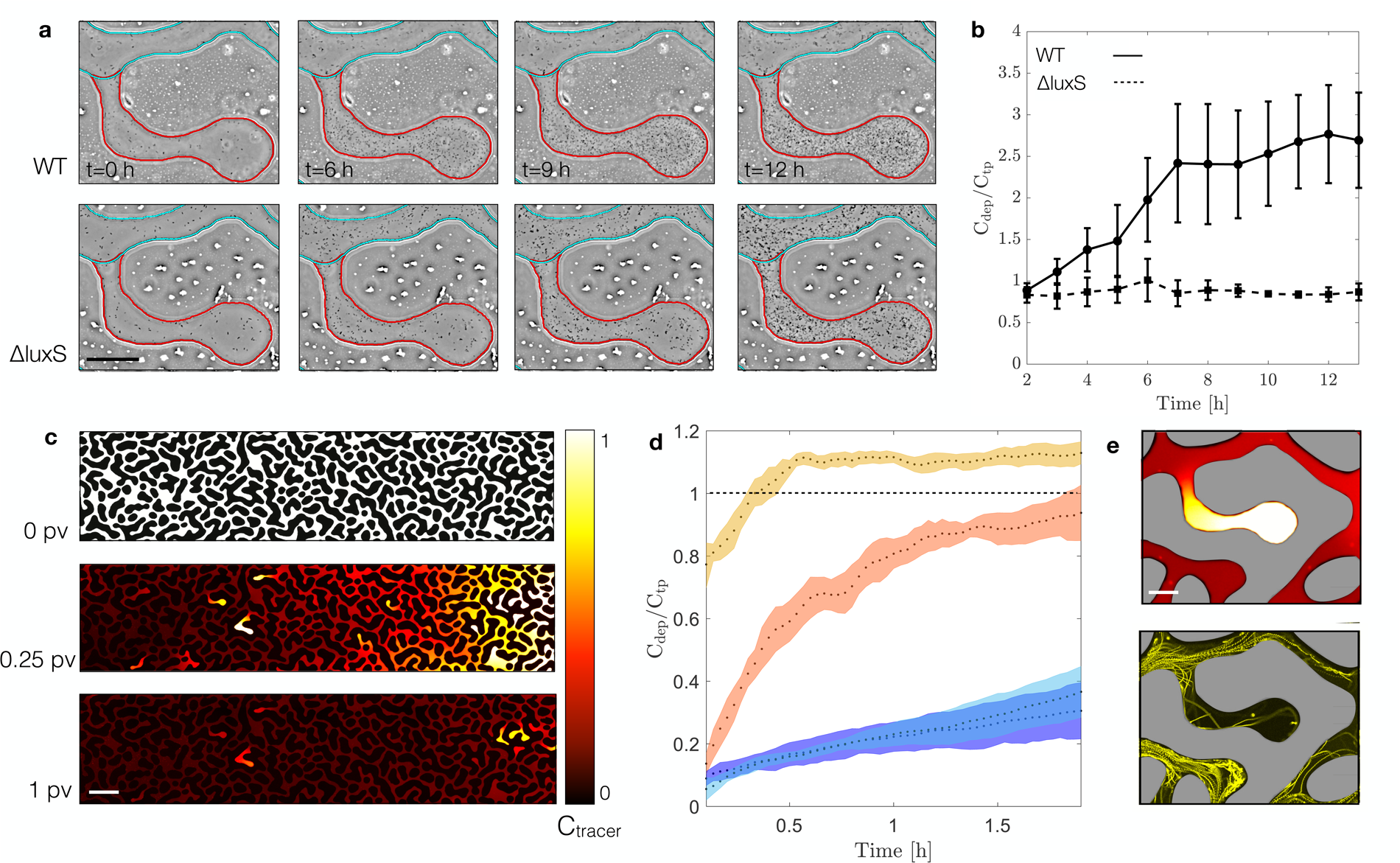
Transport mediated accumulation of *E. coli* at early times (0-12 hours). (A) Images at different times representing the bacterial abundance in a single DEP, highlighted in red, and the nearest TP, highlighted in cyan, for *E. coli* WT (top panels), and *E. coli ΔluxS* (bottom panels). Scale bar 50 µm. (B) Retention curves representing the ratio between the concentration of bacteria in the dead-end pores (C_dep_) and their concentration in the transmitting pores (C_tp_) at different times, for *E. coli* WT (straight line), and *E. coli ΔluxS* (dashed line), (*p* < 0.0001). (C) Normalized concentration map of fluorescein sodium salt at different times while displaced by de-ionized water. Scale bar 200 µm. (D) Retention curve representing the ratio between the concentration of bacteria in the dead-end pores (C_dep_) and their concentration in the transmitting pores (C_tp_), at different times. Experiments performed with *E. coli* WT (dark orange) and *E. coli Δflim* (dark blue), in absence of a chemoattractant. Experiments performed with *E. coli* WT (light orange) and *E. coli Δflim* (light blue), in presence of a chemoattractant. (E) Gradient of the fluorescent tracer (top panel) and trajectories of *E. coli* WT attracted towards a chemoattractant, after 1 PV (24 min). Scale bar 50 µm.

We examine the persistence of solute gradients within DEP-entrance and the consequent bacterial chemotaxis via two separated experiments. In the first, we qualitatively visualize the variation in intensity of an aqueous fluorescent dye (Fluorescent Sodium Salt, Merck) solution, initially saturating the same microfluidics and then displaced by clean deionized water. Fluorescent dye concentration gradients at DEPs entrance can be detected even after eluting about 1 pore volume which corresponds to 24 minutes (Fig. 3c). Thus, the gradients of this passive tracer persist for much longer than the diffusive time scale over the average DEP depth *L=0.2* mm, which is Τ_d_ = L^2^/D about 1 minute (D = 0.0006 mm^2^/s is the measured fluorescent dye diffusion coefficient [44]). This strong persistence results from the coupling between the flow kinematics [45,46] and molecular diffusion [44]. In our experiments with bacteria, local gradients across DEPs (where AI-2 is continuously produced by resident cells) and TPs (where AI-2 is continuously removed by advection) can persist much longer than for the passive fluorescent tracer.

In the second experiment, we examine if these small scale and persistent gradients are sufficient to trigger the chemotactic motion by *E. coli* cells. We saturate the microfluidic device with a solution of motility buffer supplemented with L-serine (10 mM), an amino acid toward which *E. coli* is chemotactically attracted via the same chemoreceptor (Tsr) used to sense AI-2 [30]. Next, we displace the resident solution by a sharp front injection of a mixture of the motile WT and the non-motile *ΔfliM* cells (1:1 ratio), suspended in the motility buffer without chemoattractant. We use L-serine instead of AI-2 to separate transport and chemotaxis from other effects triggered by the AI-2 availability, such as the induction of auto-aggregation genes [36]. The non-motile strain *ΔfliM* functions as a control, in this experiment. In presence of the chemoattractant L-serine, after eluting one pore volume (24 minutes) the motile cells invade the DEPs 40 % more (CDep / CTP = 1), than in the absence of the chemoattractant (CDep / CTP = 0.6). Their abundance quickly reaches a plateau as shown by the retention curve, quantified as the ratio between bacterial availability per pore feature (Fig. 3d). These two experiments show that the medium structure promotes gradients of chemoattractant along the DEPs that can be sensed by bacteria, which swim towards their higher concentration (Fig. 3e). These results justify that motile bacteria continuously accumulate within DEPs during the injection of sterile nutrient solution and promote cells encounter events observed during the subsequent times.

### Intermediate times - Transport and chemotaxis determine *E. coli* local architectural differentiation into sessile colonies and suspended cell aggregates

At times larger than 12 hours, our experiments show that cells within the DEPs start forming aggregates. Similar observations in earlier studies have been attributed to chemotaxis towards AI-2 that promotes cell-to-cell aggregation in *E. coli,* a process mediated by cell collision and aggregating factors as outer membrane adhesins, fimbriae, pili or curli fibers [30,31]. The current understanding of this mechanism is limited to relatively simple geometries without flow, where gradients of self-produced signaling AI-2 molecules persist for a short period (i.e. tens of minutes[31]), before diffusion mixes AI-2, dissipating its gradients with subsequent loss of chemotaxis. This results in disaggregation, because motile cells detaching from aggregates are in higher in number than those that join them [31]. We argue that in complex landscapes, as in our experiments, the persistence of AI-2 gradients is sustained by the combined effect of advection, constantly removing AI-2 molecules from TP, and production of AI-2 from cells trapped within DEPs. Therefore, motility towards such gradients induces aggregates formation without exhibiting disaggregation, up to about 30 hours, as visible in Fig. 4a. We further show that not only biomass accumulates differently across the two strains, but also the architectural differentiation is pore-class dependent. In the TPs, the WT strain forms a homogenous layer of cells and reaches its carrying capacity around 30 hours, while in the DEPs cell aggregates continue to increase in size until the entire DEP space is occupied (Fig. 4a,b). In contrast, the Δ*luxS* strain forms a sessile layer of biofilm, which is architecturally the same in TP and DEP (Fig. 4a,b). This is captured in the aggregate-averaged mass distribution (PDF) within the DEP that stays nearly constant for Δ*luxS* but keeps shifting towards larger values for WT, shown in Fig 4b. This implies that the increase in aggregate size is a combination of binary cell division and recruitment of new cells. Interestingly, increasing the concentration of glucose by 10 folds within the injected solution did not affect biomass accumulation of the WT in DEP. We explain this arguing that nutrient consumption by resident cells could not be balanced by diffusive transport of the resource in DEP. Thus, only cells in the TP exhibited a higher carrying capacity (Supplementary Fig. S2), as the injected solution carries fresh nutrient. Eventually, due to chemotaxis and cells aggregation, the pore space becomes crowded and the overall biomass accumulation slows down.

**Figure 4.**
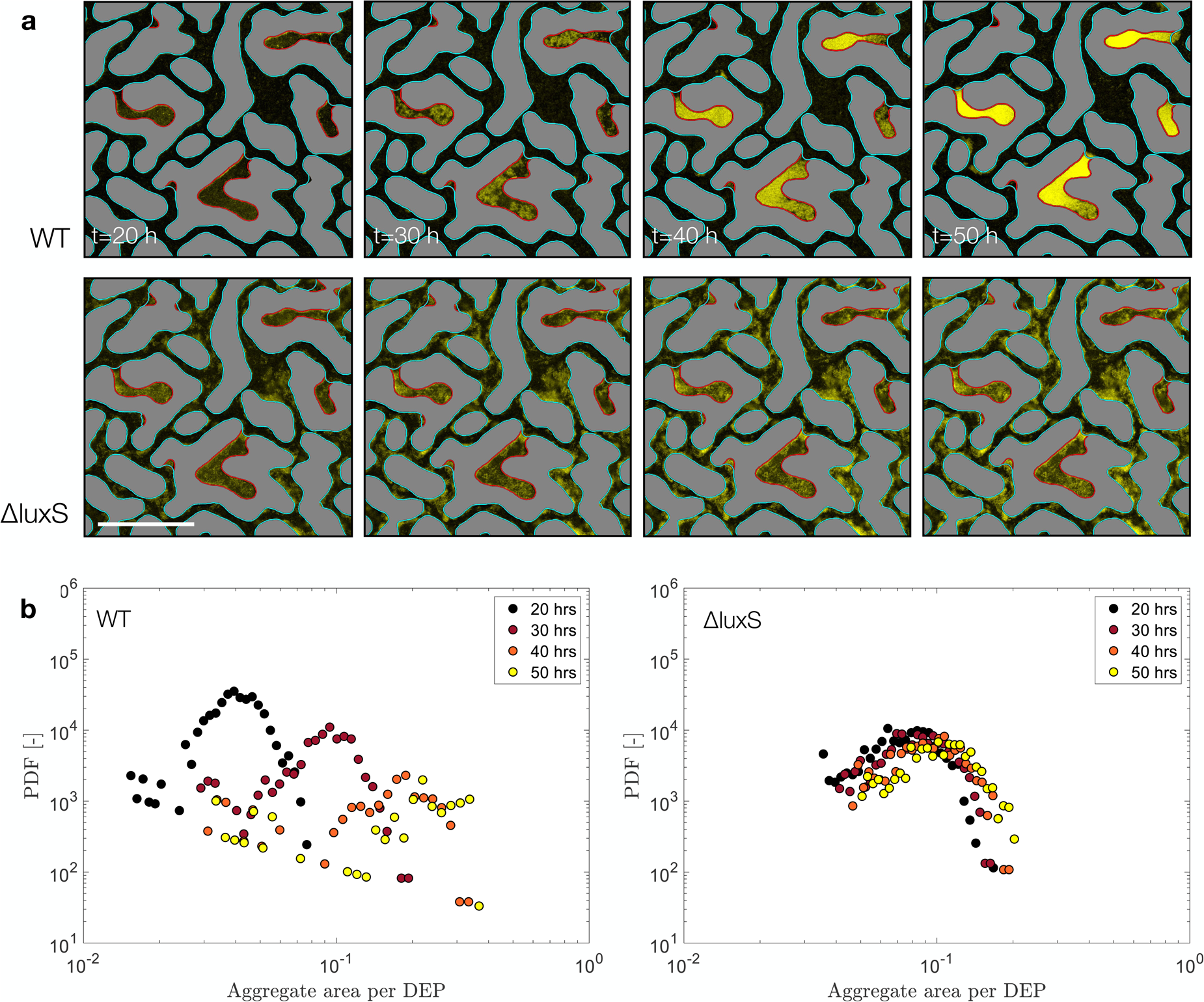
Accumulation dynamics of *E. coli* in presence or absence of AI-2, cells are shown in yellow over a dark background. (A) Images showing qualitatively the colony growth of *E. coli* WT (top panels), and *E. coli ΔluxS* (bottom panels) at different times. Scale bar 200 µm. (B) Probability density function (PDF) of individual colony biomass per colony surface, at different times, for *E. coli* WT (left panel), (*p* < 0.0001), and *E. coli ΔluxS* (right panel), (*p* > 0.1).

### Late times - Nutrients limitation and Quorum Sensing control biofilm formation in DEPs

After 30 h, WT biomass accumulation in the TPs slows down and almost stops (see Fig. 2b). In contrast, in the DEPs the biomass accumulates faster than in the intermediate phase (12-30 hours), until the growing biofilm extrudes from an individual DEP into the nearest TP (Fig. 5a). We, therefore, hypothesize that around 30 h the high levels of AI-2 induce QS and activate a gene expression that controls biofilm phenotype, a QS regulation known to occur in *E. coli* [36,47]. In this regard, the gene *lsrR* plays an important role for *E. coli* by directly controlling AI-2 uptake and regulating the expression of several important biofilm-related genes such as *wza*, involved in the synthesis of colanic acid a building block of EPS, and *flu*, which encodes the proteins that mediates cell-to-cell aggregation [32,33].

**Figure 5.**
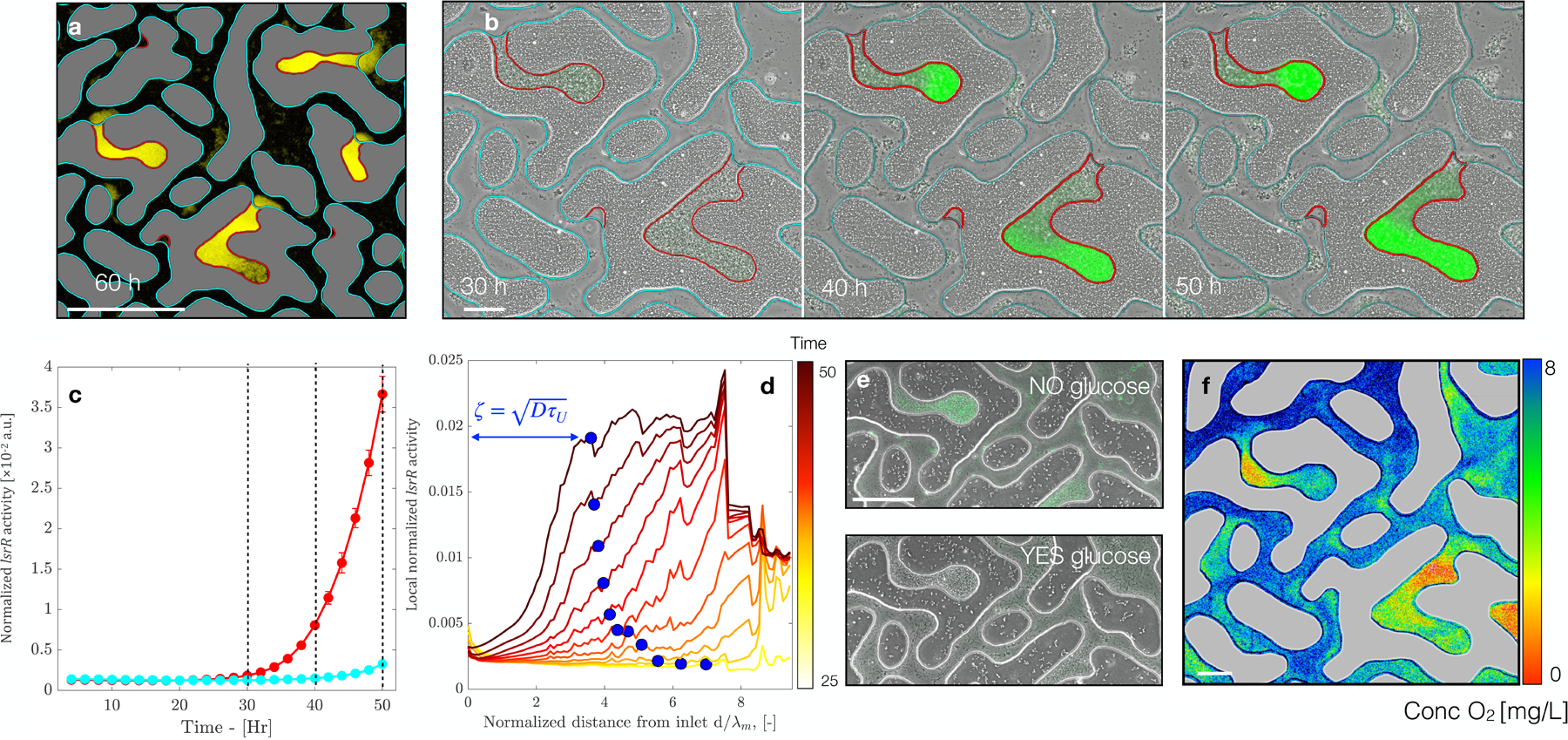
QS induction of biofilm formation in DEPs. (A) Biomass extruding out of the DEPs after 60 hours. Scale bar 200 µm. (B) Images showing the biomass accumulation in one DEP (phase contrast) and the P*_lsrR_* activity (green superimposed color) of *E. coli* WT at different times. Scale bar 50 µm. (C) Quantification of the P*_lsrR_* activity, normalized by biomass abundance for *E. coli* WT, (*p* < 0.0001) in TP, cyan, and DEP, red. (D) Averaged normalized P*_lsrR_* activity along DEP length, from length to dark increasing time. The glucose limitation length scale, ζ, is represented with a blue circle. (E) Images representing the biomass accumulation in one DEP superposed to the P*_lsrR_* activity (green) of *E. coli* WT, in LB 0.1 no glucose (top panel), and LB 0.1 amended with 5mM of glucose (bottom panel). Scale bar 100 µm. (F) Map of the oxygen concentration resulting from the coupling of advective-diffusive transport and *E. coli* WT uptake after 50 hours of experiment. Scale bar 50 µm.

To test if the observed biomass accumulation is the result of QS, we estimate when and where *E. coli* cells start to internalize AI-2, and if there is a temporal agreement with the quantified biomass accumulation dynamics. To this end, we perform a set of three independent experiments where cells of *E. coli* are equipped with a *lsrR* gene reporter (pMSs201-P_lsrR_-gfp) in which the *lsrR* promoter (*P_lsrR_*) is fused to a gene encoding Green Fluorescent Protein (GFP). As in the experiment with the WT, cells of the reporter strain *P_lsrR_*, previously injected in the microfluidic device, are exposed to the same constant injection of nutrient sterile solution (a M9 medium supplemented with glucose 5mM). For the first 25 hours the reporter activity is off (not detected). However, when concentration of cells in DEP is high (around 30h) the overall reporter activity normalized per unit of biomass, shows an increasing expression of the *lsrR* gene in the DEPs, which does not occur in the TPs (as shown in Fig. 5b-c).

Glucose metabolism, however, directly modulates QS: in presence of the sugar, catabolite repression inhibits AI-2 uptake, therefore hindering the gene expression that controls biofilm production [34,39,40]. This is because the phosphotransferase sugar transport system (PTS) member (HPr) exists in the unphosphorylated form and prevents lsr system to import AI-2. As glucose is depleted, HPr is mostly converted in p-HPr, and lsr inhibition is released [39]. We argue that, despite AI-2 concentration rises above the quorum threshold, AI-2 uptake and eventually the phenotypes coordinated by QS is expressed only when glucose is depleted in the DEPs. Therefore, we further measure the reporter activity along the depth of all DEP (defined as the GFP signal of the reporter averaged along the skeleton of each individual DEP structure; see Methods). Figure 5d (solid lines, color coded by time) shows the reporter activity per unit mass along each DEP and averaged over all DEPs: the signal is strong deep down each DEP, and it becomes progressively weaker towards the entrance (at the juncture with the adjacent TP). The transition between high to low QS reporter activity takes place at a distance ζ away from the DEP-TP juncture, at which the normalized *lsrR* activity indicates that AI-2 is imported and therefore the glucose availability must be limited. We assume that this length scale ζ represents the limit of the glucose depleted zone within the DEP: it results from the competition between glucose consumption by microbial cells and its diffusion from the nearest TP through the biofilm itself. The glucose consumption time τ_U_ can be estimated knowing the DEP-averaged biomass estimated from each image (see Methods). In Fig. 5d (blue circles), we represent the modeled glucose limitation length, 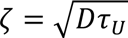, where τ_U_ is estimated from the average DEP biomass observed and assuming a constant glucose concentration of 5 mMol at the DEP-TP juncture (which is the injected concentration). To exclude that AI-2 is not taken up in presence of glucose, we perform two more control experiments in which cells of *E. coli* reporter strain *PlsrR* are exposed to a constant flow of LB (10% dilution) with and without 5mM glucose. Despite the two conditions result in a similar biomass accumulation, we observe contrasting results in the reporter activity. In presence of glucose, the reporter activity remains off at all times, while without glucose and in presence of LB, the reporter shows higher activity in the DEPs than in the TP (Fig. 5e and Supplementary Fig. S3).

## Discussion

The mammalian gut commensal *E. coli* coordinates population behaviors via secreting, sensing, and up-taking the chemoattractant and QS signaling molecule, AI-2 [40]. In a spatially complex micro-environment, the local availability of dissolved AI-2 is controlled by the coupling of fluid flow in advection dominated zones connecting areas of fluid stagnation [41,49]. In this work, we use microfluidic experiments and automated time-lapse video-microscopy to explore the role of different structures, Transmitting and Dead-End pores, and the consequent flow heterogeneity on bacterial colonization of a system that mimics the spatial complexities in the gut [23]. We show that flow heterogeneity translates into persistent gradients of signaling molecules and carbon resources (here, glucose) with consequences on the colonization and spatial organization of *E. coli.* Specifically, we distinguish different mechanisms controlling the colonization of *E. coli*: a) chemotaxis driven by gradients of self-produced chemoattractant AI-2, b) aggregation mediated by cells collision, and c) AI-2 uptake which regulates biofilm phenotype formation via QS. Despite the overall amount of biomass growth for *E. coli* WT and Δ*luxS* mutant (that do not produce AI-2) are similar after 48 hours, their spatial distribution and growth dynamics differ significantly.

We show the *E. coli* production of AI-2 and its accumulation within DEPs promote chemotactic migration towards and cells accumulation within the DEPs that, at later times, triggers QS and enhanced biofilm formation. This pore-specific colonization does not occur for a Δ*luxS* mutant; therefore this phenomenon depends on the ability of the WT strain to produce and sense AI-2 gradients. Taken together, these findings show that chemotaxis and QS control the spatial segregation of bacterial strains based on the strain-specific response to AI-2. This is an indication of an active role of AI-2 secretion and sensing in initiating ecological niche segregation [50,51], and therefore stable co-existence of different *E. coli* strains in spatially complex environments. This is supported by results of competitive experiments with *E. coli* in murine gut, where strains characterized by different AI-2 detection capability (WT and Δ*lsrB*) coexist by occupying different niches [52].

While chemotaxis is energetically costly, it entails several benefits such as efficient nutrients acquisition, avoidance of harmful substance, formation of biofilms and colonization of new environments [53]. We show that chemotaxis allows *E. coli* to colonize cavities such as the interspace between villi and crypts, where they likely outcompete local invaders due to their higher local population. Additionally, if a bacterial strain is able to detect AI-2 more efficiently than other strains, it may gain a competitive advantage, as shown by comparing a WT with a non-chemotactic mutant (Δ*cheY*) [52]. Concentration and quality of carbon availability may also influence the bacterial chemotactic response. In *E. coli*, the expression of motility genes is inversely proportional to the growth rate achieved on different carbon sources [54]. In most conditions, glucose is the best carbon source for *E. coli*, and for this study, we chose a concentration comparable with that found in mammalian guts (5-50 mM) [43]. In presence of other resources, the active role of motility could therefore be more or less relevant, and resulting in alternative colonization pattern.

Chemotaxis toward AI-2 mediates cell-to-cell aggregation, until diffusion mixes the gradient [30,31]. In our model system, the heterogeneous flow and AI-2 production sustain AI-2 gradients over times comparable with the ones allowing the establishments of a mature biofilm. The cell accumulation driven by chemotaxis along the gradients towards DEPs for such long times promotes the formation of bacterial aggregates, whose formation is mediated by cell-to-cell collision in the confined space of a DEP. Notably, these formations do not disaggregate after 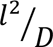 when diffusion is supposed to have homogenized AI-2 concentration over the aggregate size *l*, therefore hindering chemotaxis as previously reported [31]. Here, we show that the medium structural heterogeneity together with fluid flow sustain aggregates formation over tens of hours making DEPs hotspots for microbial accumulation and aggregation.

At later times, we observe that biomass accumulation within the confined space of DEPs induces QS that mediates enhanced biomass growth leading to the transition towards a sessile biofilm life style. Furthermore, using a fluorescent reporter of *lsrR* expression, a gene involved in the uptake of AI-2 and controlling biofilm phenotype [47,55], we conclude that QS and the transition towards sessile biofilm life style happens as a response to resources limitation. We show that AI-2 uptake begins from the end of each DEPs, presumably only once glucose is depleted and diffusion from the nearest TP is not sufficient enough to provide more glucose. To sustain this conclusion, we computed the diffusive length scale ζ that glucose travels along the DEP before becoming completely depleted by cells uptake. Our model shows that this length scale well predicts the fluorescent signal front emitted by the *lsrR* along DEP.

Evidently, another carbon source is needed by *E. coli,* to sustain the biofilm formation that QS coordinates once glucose is depleted. It is known that glucose is partially catabolized and excreted as acetate by *E. coli*, a process known as acetate overflow metabolism [56]. Only once acetate-producing carbon sources are depleted (glucose, herein), *E. coli* can scavenge for the previously produced acetate via acetyl-coA synthetase (*acs)*. This mechanism, called acetate-switch [57], has been reported for *E. coli* growing within confined environment [58]. Interestingly, in *Vibrio fischeri,* a squid symbiont, this metabolic switch has been recognized to be coordinated by QS via controlling *acs* [59,60]. This synthetase provides an overall advantage in colonizing the crypts of the squid light-organs, compared to *acs* deficient strains [59,60]. To what extent the two processes (QS and acetate-switch) are coupled in *E. coli*, remains to be understood.

*E. coli* cells uptake AI-2, however they do not use it as an energetic resource [61]. This metabolic expense has been shown to return a colonization advantage by interfering with neighboring cells [33]. We propose here another advantage: QS is beneficial for bacteria to mechanically escape unfavorable condition, via enhanced production of biomass. To test this hypothesis, we equipped the microfluidic device with transparent oxygen-sensors [62] to estimate the availability of dissolved oxygen within the two pore classes. As expected, after 30 hours the dissolved oxygen concentration is lower within the DEPs due to its consumption via cell-respiration and reduced renewal due to the tortuous structure of a DEP non directly accessible to flow (Fig. 5f). Hence, the excess of biomass extrudes from DEPs to access zones with higher oxygen and glucose concentration, both continuously flowing within the TPs. This is corroborated by biofilms models that incorporate reaction-diffusion effects, where it has been shown that biomass production pushes daughter cells towards a more oxygen rich environment [63]. Our findings support the hypothesis that bacterial cells within confined environments invest energy via QS (biomass production) to turn resource depleted zones into hot-spots of intense activity and mechanically escape unfavorable conditions.

To conclude, understanding how bacteria colonize complex and heterogeneous environments, such as the gut, is essential for comprehending host-microbe interactions and their impact on health and disease. We show how the spatial structure and flow heterogeneity of these environments play a crucial role in shaping bacterial behavior and colonization, particularly through chemotaxis and quorum sensing. This is of great relevance, as AI-2 is a universal inter-species communication system [33], and different bacterial species can produce and respond to it, modulating gut microbiota composition [64]. We showed here how persistent chemotactic gradients and QS may affect bacterial behavior in systems characterized by complex structures for an isolate strain and, thus likely, also for multispecies gut microbial communities.

## Methods

### Bacterial strains and growth condition

All experiments were performed using *E. coli* MG1655 and its derivative strains. We used the motile wild type *E. coli* MG1655 pME6012-P_tac_-gfp (WT), the non-flagellated mutant *E. coli* MG1655 pME6012-P_tac_-mcherry (*ΔfliM*), the mutant lacking the AI-2 synthase gene luxS, *E. coli* MG1655 pME6012-P_tac_-gfp (*ΔluxS*), and *E. coli* MG1655 pMSs201-P*_lsrR_*-gfp. We grew the strains in 4 mL of lysogeny broth Miller (LB) with 25 µg/mL of kanamycin at 37°C incubation temperature, while shaking at 180 rpm. We diluted the overnight culture at 1:100 in 4 mL of LB, and incubated under the same conditions until the exponential growth phase (∼3 hours) was reached. Next, we centrifuged an aliquot (2300 g, 5 min), removed the supernatant, and resuspended the pellet in the motility buffer. The motility buffer consisted of 10 mM potassium phosphate, 0.1 mM EDTA, 10 mM lactate, 1 mM methionine, pH 7.0 [65]. We performed the bacterial growth experiments in M9 minimal medium, supplemented with 2 mM MgSO4, 0.1 mM CaCl2, and 5 mM or 50 mM glucose.

### Microfluidic devices

We designed the porous micromodel exploiting the method of solid-state dewetting, which results in a surface morphology exhibiting spinodal-like structure and a disordered hyperuniform character described in [23]. We printed the micromodel geometry into a silicon wafer via soft lithography, depositing a layer of SU-8 2150 (MicroChem Corp., Newton, MA) with controlled thickness (0.05 mm) via spin-coating. The wafer acts as a mold for liquid polydimethylsiloxane mixed with 10% by weight with its own curing agent (PDMS; Sylgard 184 Silicone Elastomer Kit, Dow Corning, Midland, MI). After solidification, we plasma-sealed the microfluidic device onto 25 mm × 75 mm glass slides.

### Microfluidic experiments

We performed the microfluidic experiments within a constant temperature microscope incubator (OKOlab) maintained at 37°C. First, we saturated the microfluidic device with a motility buffer and subsequently injected the bacterial suspensions via two inlet-ports operated on a 3-way-valve system. This system allowed to generate a sharp front of suspended bacteria at the chip inlet, as described in [23]. We imposed a constant flow rate (*Q*), of 0.1 μL/min with a syringe pump (PHD-ULTRA, Harvard Apparatus) eluting one pore volume (PV, i.e. the volume of the entire porous channel) every 24 min. After 5 PV, we switched the inflow from the bacterial suspension to the sterile nutrient medium keeping the flow rate same and recorded the biomass accumulation over 2 days.

### Porous medium characterization

At the end of each experiment, we created a “mask” image to characterize the porous structure. To this end, we saturated the porous medium with a florescent dye (fluorescein sodium salt, from Merck, visible through a GFP filter cube) and recorded a large image of the entire structure. After thresholding, binarizing the mask image (pore-space: 1; solid-space: 0), we geometrically discretized the two pore classes using the maximum inscribed circle method described in [23]. From the binary image we obtained then the skeleton, a 1-pixel width representation of the pore space (Supplementary Fig. S4a). We assigned a segregation index (**ζ**) to the pore-regions based on the number of neighboring grains. Hence, a segregation index (**ζ**) of 1 isolated the dead-end pores from the transmitting pores with **ζ** > 1. This characterization allowed to generate a “mapped mask”, where grains have been attributed a pixel value of 0, dead-end pores a pixel value of 1, and transmitting pore a pixel value of 2 (Fig. 1c, gray, red and light blue, respectively). The medium is quite homogeneous, such that the statistical distribution of *λ* is narrow and has a strong peak close to the mean pore-size *λ_m_* = 0.04 mm (Supplementary Fig. S4b).

### Microscopy

We performed time-lapse imaging with an automated transmission light microscope (Eclipse Ti2, Nikon) equipped with a CMOS camera (Hamamatsu ORCA flash 4.0, 16-bit, 6.5 μm per pixel) and controlled by the software Elements. This integrated system allows for automatic capture of large image (stitching 3×9 individual pictures) time series for the entire porous domain, with multichannel optical configuration. We collected individual pictures (2048×2048) with a Nikon objective 10X magnification (0.65 µm/pixel) for each time series in bright field, phase contrast and fluorescence optical configurations. For the fluorescence optical configuration, we used Nikon GFP HQ and mCherry HQ filters coupled with a Spectra X light engine.

### Bacteria accumulation analysis

We quantified pore-space accumulation of bacteria based on the detected pixel-intensity attenuation from the time-series of large images captured using the bright field configuration, as in [10]. First, we removed the background noise by subtracting from each large image the one taken at the beginning of the experiment before cells injection. Next, we multiplied each large image with a binary mask to isolate the accumulations within the transmitting pores. We processed each dead-end pore independently and identified the boundaries of individual clusters of accumulated biomass using an intensity threshold (based on mean of regional maximum intensities). We computed the area of individual clusters to compute the local concentration distribution by taking the sum of all intensities within each cluster normalized by the respective cluster area (see Fig. 4b).

### Reporter analysis

We followed the same protocol for the images acquired in bright field but in fluorescence configuration (GFP filter cube), with fluorescence intensity being a proxy of the *lsrR* activity. We discretized between DEP and TP, and then normalized the fluorescence intensity by the intensity of the biomass acquired in bright field. We further measured local biomass abundance and reporter activity along the depth of all DEP, by averaging gfp and light intensity signals within the maximum inscribable disks centered along the skeleton of the individual DEP structure, every 10 pixels (Supplementary Fig. S5).

### Particle tracking

To measure motility speed of individual *E. coli* cells (both WT and Δlux), we performed an independent experiment to track trajectories of bacteria under the no-flow condition. After loading the bacteria suspension (in motility buffer) into the microfluidic channel, we insulated the inlet/outlet ports. We captured time-series images using phase-contrast configuration at a magnification of 15X and acquisition rate of 100 frames/sec with an exposure time of 2 ms for a total duration of 40 s. To reconstruct the trajectories of individual bacterial cells, we follow the same method described in [7]. The velocity PDF based on approximately 7500 trajectories for both bacteria strains are shown in Supplementary Fig. S1.

### Numerical simulation of flow velocity

We performed a stationary creeping flow simulation using COMSOL Multiphysics over a smaller microfluidic domain of 9.6 mm × 3.6 mm with the same structure, as in [23]. The computational resolution is high enough to ensure a divergence free velocity field.

### Consumption-diffusion of glucose in DEP

The limit ζ of the glucose depleted zone within the DEP results from the competition between glucose consumption by microbial cells and its diffusion from the nearest TP through the biofilm itself. We estimate the glucose consumption time τ*_U_* by the cells within a DEP as the uptake rate *U_1c_*: per dry mass of *E. coli* (10 mMol per second per g of dry mass of bacteria [66]) multiplied by the DEP biomass, *M_dc_*. The latter is estimated by assessing, first, the average number of cells in a DEP from each image. Then, the mass of cells in DEP is given by the product of their estimated number and the dry mass of an individual cell (e.g.[67]): finally, the average glucose uptake rate in the DEPs is *U_T_ = U_1c_M_dc_*. Then, assuming a constant glucose concentration *c_0_* of 5 mMol at the DEP-TP juncture (which is the glucose concentration in the injected solution), we get τ*_U_* = *c*_0_/*U*_T_. Thus, this transition between high and low glucose is supposed to superpose to the measured reporter (*P_lsrr_*) signal along the DEP length.

### Oxygen measurement

We optically quantified the dissolved oxygen within the medium pores by means of transparent planar optode sensors built within the microfluidics [62]. The planar sensor consists of a thin layer (< 3 µm) of a solid polymer matrix containing two fluorescent dyes. The first is quenched by molecular oxygen whereas the second dye provides a reference signal. We independently imaged the local fluorescent signal intensity of the two dyes by means of two customized filter cubes (450/650 nm Ex/Em and 450/520 nm Ex/Em). The oxygen map is then derived from the ratio between the signal of the two dyes.

### Statistical analysis

One-way analysis of variance (ANOVA) was carried out when two or more treatment groups were compared, and two-sample t-test when comparing two groups. Statistical analysis was performed using the software Matlab. P values <0.05 were considered to indicate statistical significance.

## Author Contributions

D.S., A.D.B. and P.d.A. designed the research, M.B. provided the de-wetted HPS samples, D.S. performed experiments, A.D.B. performed numerical simulations, D.S., A.D.B. F.E. and P.d.A. analyzed the data, V.S. prepared the reporter strains and all authors wrote the manuscript. D.S. and A.D.B. share the first authorship.

## Acknowledgements

We thank Jan Roelof van der Meer and Thomas Keith Wood for providing the strains. We thank Sergey Borizov for providing the oxygen optode sensors used in this study.

