## Supplementary Information for "Spatial structure, chemotaxis and quorum sensing shape biomass accumulation in complex systems"

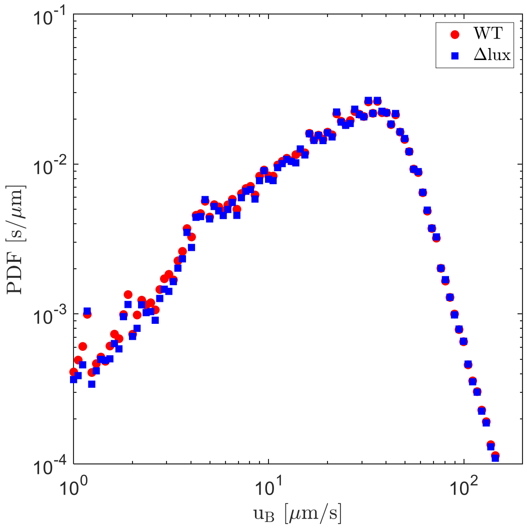


**Figure S1.** Distribution of the swimming velocities of WT (red) and Δ*luxS* (blue) derived from the statistical analysis of particle tracking performed in absence of flow ($p > 0.1$).

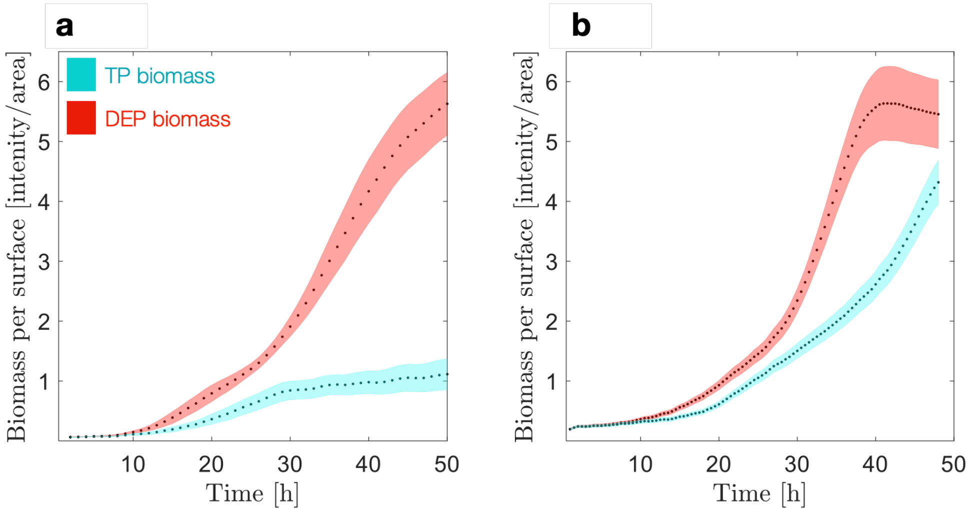


**Figure S2.** Impact of glucose concentration on *E. coli* WT carrying capacity. Comparison of biomass concentration per pore type, for (A) experiments performed at glucose concentration of 5 mM, and (B) experiments performed at glucose concentration of 50 mM.

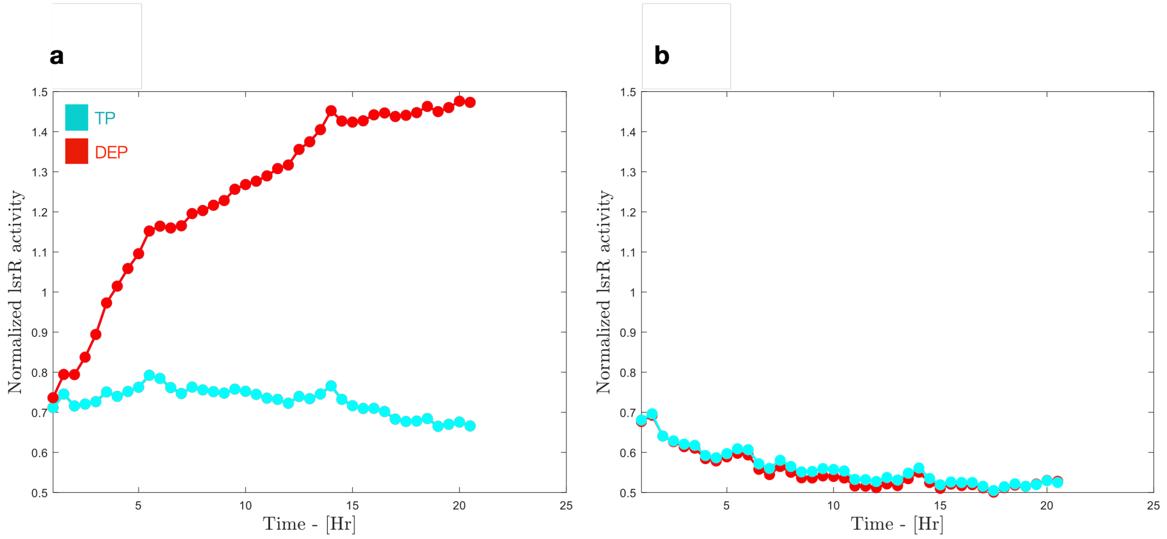


**Figure S3.** Impact of glucose concentration on *E. coli lsrR* activity. Comparison of normalized *lsrR* activity per pore type, red for dead-end pores, cyan for transmitting pores. (A) Experiments performed in Lysogeny Broth diluted in distilled water 10 times (LB 0.1) without glucose, ($p < 0.0001$). (B) Experiments performed at with LB 0.1 and glucose 5 mM ($p > 0.1$).

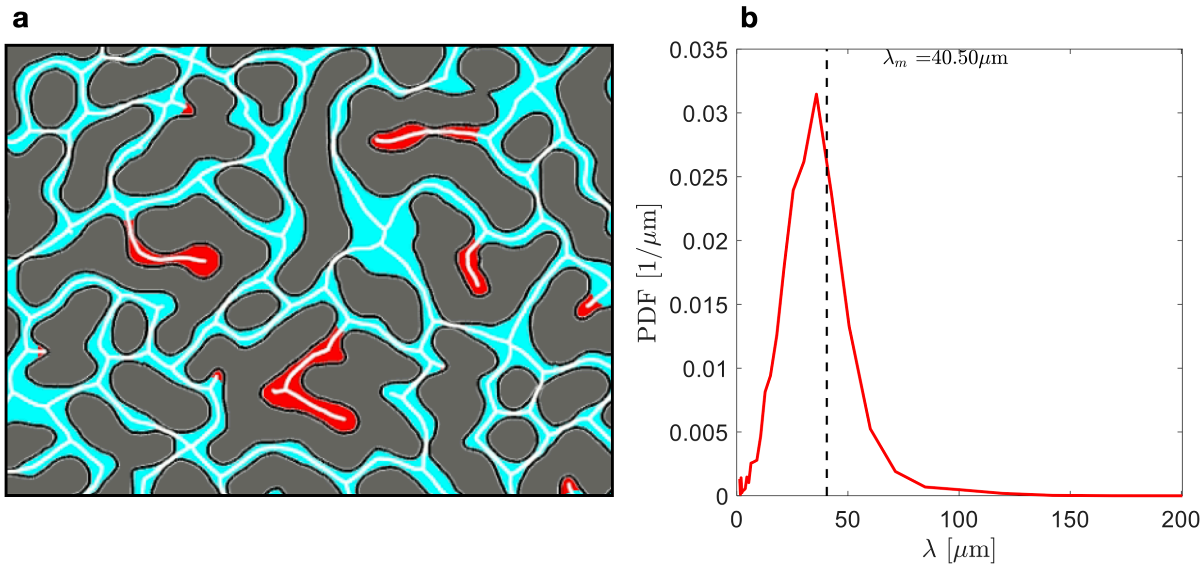


**Figure S4.** Pore throat characterization. (A) Representation of the binarized map where TP are colored in cyan, DEP in red, and skeleton in white. (B) Distribution of pore throats measured along the skeleton of the system, dashed line indicates average pore throat size, $\lambda_m$ = 40 µm.

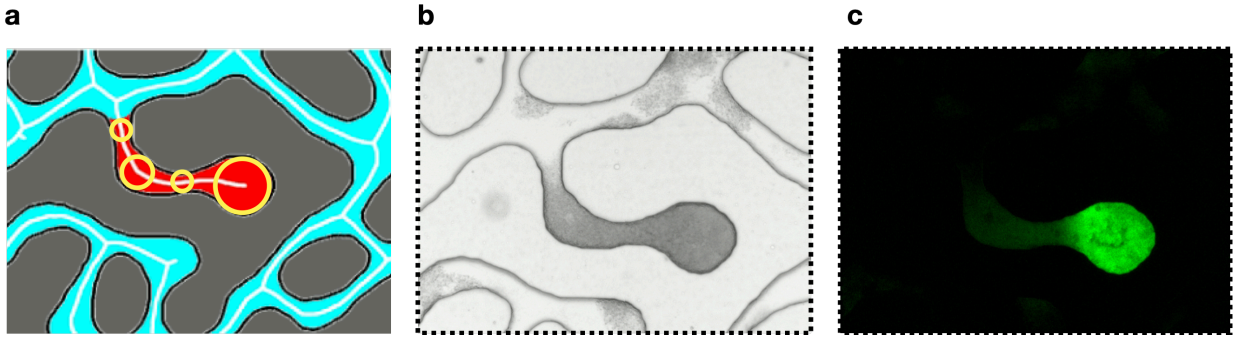


**Figure S5.** Pore scale measurement of biomass and *lsrR* activity. (A) Image intensities have been averaged over a disk of radius *r* centered along the skeleton of each individual DEP structure, every 10 pixels. In figure have been plotted only 4 representative circles along the skeleton. This allowed to quantify (B) the biomass accumulation from the bright field images, and in the same way to estimate (C) the *lsrR* activity from the reporter fluorescence signal.
